# Myristoyl’s dual role in allosterically regulating and localizing Abl kinase

**DOI:** 10.1101/2022.12.20.521177

**Authors:** Svenja de Buhr, Frauke Gräter

## Abstract

c-Abl kinase, a key signalling hub in many biological processes ranging from cell development to proliferation, is tightly regulated by two inhibitory Src homology domains. An N-terminal myristoyl-modification can bind to a hydrophobic pocket in the kinase C-lobe, which stabilizes the auto-inhibitory assembly. Activation is triggered by myristoyl release. We used molecular dynamics simulations to show how both myristoyl and the Src homology domains are required to impose the full inhibitory effect on the kinase domain, and reveal the allosteric transmission pathway at residue-level resolution. Importantly, we find myristoyl insertion into a membrane to thermodynamically compete with binding to c-Abl. Myristoyl thus not only localizes the protein to the cellular membrane, but membrane attachment at the same time enhances activation of c-Abl by stabilizing its pre-activated state. Our data put forward a model in which lipidation tightly couples kinase localization and regulation, a scheme that currently appears to be unique for this non-receptor tyrosine kinase.

## Introduction

The non-receptor tyrosine kinase c-Abl, hereafter referred to as Abl, is involved in a plethora of signalling processes, including cell proliferation and survival, stress response, neuronal development and remodeling of the actin cytoskeleton (***Khatri et al., 2016***; ***Pendergast, 2002***; ***Wang, 2014***; ***Woodring et al., 2003***). Knock-out of Abl leads to developmental lethality (***Koleske et al., 1998***), and fusion with the breakpoint cluster region gene from chromosome 22 yielding the BCR-Abl oncogene de-regulates kinase activity and is the main cause of chronic myelogenous leukemia (CML) (***Greuber et al., 2013***). Given its key signalling function, tight and multi-modal regulation of Abl kinase activity is essential, and allostery has emerged as an important regulatory principle and therapeutic target for this large multi-domain kinase (***Nussinov et al., 2022***).

From N- to C-terminus, Abl consists of a flexible linker referred to as the N-cap, two Src homology (SH) domains, SH3 and SH2, the catalytic kinase domain, a disordered region containing proline-rich motifs and localization signals, and an F-actin binding domain. The assembled, auto-inhibited structure is similar to Src family kinases and the two SH domains attach to the back of the kinase domain with respect to the active site (***Nagar et al., 2003***). Isoform 1b of Abl is myristoylated at its N-terminus. The myristoyl moiety (Myr) binds to a hydrophobic pocket in the kinase domain, which induces a kink in the kinase C-terminal *a*_I_ helix. This kinked conformation of the helix enables docking of the SH2 domain to the kinase C-lobe and consequently kinase inhibition (***Hantschel et al., 2003***). Apart from this, two more characteristics contribute to stable inhibition. The PxxP motif on the linker connecting SH2 domain and kinase facilitates stable docking of the SH3 domain (***Panjarian et al., 2013***; ***Ren et al., 1993***), and the N-cap acts as a rigidifying clamp around the SH3-SH2-kinase complex (***Chen et al., 2008***). While many aspects of Abl activation have been examined, a comprehensive picture starting from this initial clamped and inhibited structure is missing.

Myr and the SH domains have to detach from the kinase domain for Abl activation. The kinase domain then undergoes several structural transitions: The *α*_C_ helix of the kinase N-lobe rotates inward, the activation (A-)loop switches from a collapsed to an extended state that allows substrate binding, the DFG-motif converts to the in-conformation capable of coordinating ATP and a Mg^2+^ ion for catalysis, and the hydrophobic regulatory (R-)spine at the hinge between N- and C-lobe assembles (***Azam et al., 2008***; ***Levinson et al., 2006***; ***Taylor et al., 2019***; ***Xie et al., 2020***). Docking of the SH2 domain to the kinase N-lobe facilitates stable transition to the active conformation (***Grebien et al., 2011***; ***Lamontanara et al., 2014***; ***Nagar et al., 2006***). Knowledge about these conformational states stems mainly from structural, i.e. rigid, data and has been complemented with hydrogen-deuterium exchange mass spectrometry or solution NMR experiments investigating the effects of ATP-competitive or allosteric inhibitors (***Iacob et al., 2009, 2011***; ***Skora et al., 2013***) or determining the energy landscape of different Abl constructs (***Saleh et al., 2017***; ***Xie et al., 2020***). Previous computational studies have been instrumental in examining the conformational transitions between active and inactive states and structural elements regulating kinase activity such as *α*_C_ helix or DFG-motif conformation (***Dölker et al., 2014***; ***Liu et al., 2022***; ***Meng et al., 2018***; ***Narayan et al., 2020***). However, surprisingly little is known about the early stages of Abl’s allosteric activation pathway directly following unbinding of its natural ligand Myr. We here aim to resolve at atomistic level how Myr allosterically regulates Abl by impacting kinase domain dynamics and auto-inhibition, and the conformation of the C-terminal *α*_I_ helix.

In contrast to the assembled state, in which the *α*_I_ helix assumes a kinked conformation, crystal structures of only the kinase domain without Myr or any other kink-inducing ligand bound display the helix in a straight conformation. This led to the common view that Myr unbinding causes straightening of the C-terminal *α*_I_ helix, impairing the SH2-kinase interface and consequently leading to the loss of inhibitory interactions from both SH domains. This is consistent with biochemical experiments showing elevated activity of Abl after Myr removal. However, a high-resolution structure of the kinase domain in complex with the SH domains, but without Myr has not been resolved. SAXS shape reconstructions and solution NMR of non-myristoylated Abl reveal the complex in the assembled state (***Badger et al., 2016***; ***Skora et al., 2013***). We here show, using equilibrium and non-equilibrium Molecular dynamics (MD) simulations, that Myr removal does not fully straighten the *α*_I_ helix but fosters a highly dynamic helix conformation, a pre-activated kinase conformation, and a loosened interface to the SH domains. We also reveal the underlying allosteric pathways at atomistic detail.

Having established a key regulatory capacity of Myr, we additionally asked how Myr binding to Abl is controlled. Our MD simulations of Myr-unbinding from Abl versus a membrane suggest that Myr unbinding and kinase pre-activation can be promoted by insertion of Myr into the cellular membrane, as Abl and a lipid bilayer can thermodynamically compete for Myr binding. Taken together, our computational study suggests a dual role of Myr as allosteric inhibitor and membrane anchor, putting forward the intriguing possibility of a direct crosstalk between Abl kinase activity and membrane localization.

## Results

### Effect of Myr on *α*_I_ helix and kinase dynamics

We first set out to explore the effect of Myr binding on the overall dynamics of the Abl kinase domain using equilibrium MD simulations. To this end, we simulated the SH3-SH2-kinase complex and systems of only the kinase with the *α*_I_ helix in a kinked or a straight conformation (Fig. 1A). All models included ATP and were simulated both with or without Myr. During the 15 µs cumulative simulation time (30 × 500 ns) per model we did not observe spontaneous straightening of the *α*_I_ helix in the absence of Myr. The straight *α*_I_ helix on the other hand unfolds regardless of whether Myr is present or not (Fig. 1B, Appendix Fig. 1). This is not unexpected since the hydrophobic residues I521, V525 and L529 (we use 1b isoform numbering throughout this work), that interact with Myr in the kinked *α*_I_ conformation, are solvent exposed in the straight conformation. The low stability of a straight helix is consistent with crystal structures of the kinase domain, in which the folded or resolved part of the helix mostly ends before or around the position of the first hydrophobic residue, while the more C-terminal residues are not resolved. Longer helices are often stabilized by another protein copy packed against it in the the crystal lattice and can therefore be regarded as a crystallization artifact (Appendix Fig. 2). Importantly, the kinked *α*_I_ helix remains folded in the absence of Myr but is strongly destabilized, as can be seen by an increased RMSF of the second half of the helix after the kink both in simulations of a single kinase domain and even, albeit to a lesser extent, in presence of the SH-domains, which has also been observed by ***Liu et al. (2022***).

**Figure 1.**
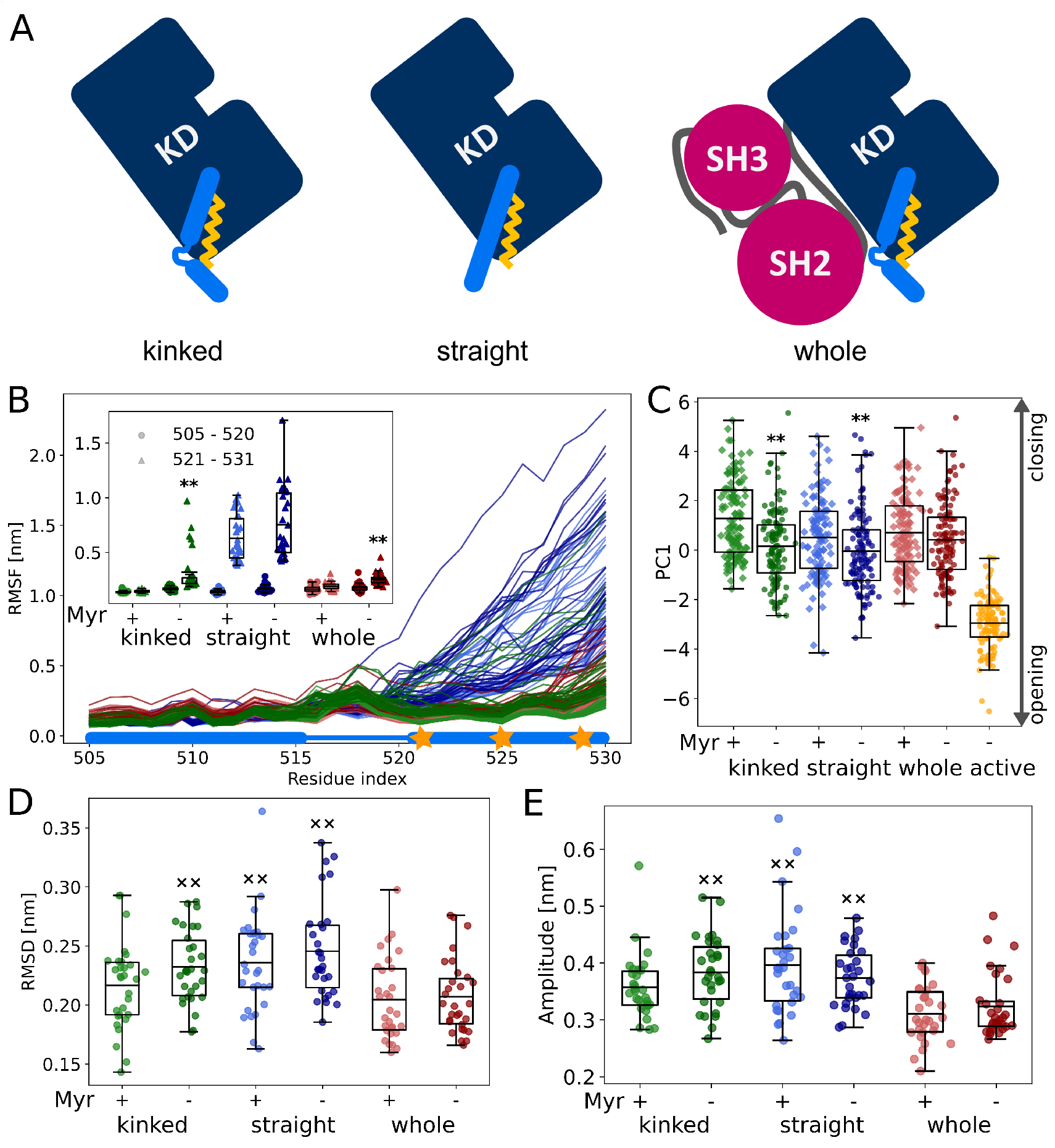
Influence of Myr on kinase domain dynamics. (A) Overview of simulated Abl models. Models referred to as ‘whole’ contain the kinase (KD, dark blue) and SH domains (magenta). The kinase was simulated with a kinked or straight *α*_I_ helix (light blue) conformation. All models were simulated in the absence or presence of Myr (yellow). (B) RMSF of the *α*_I_ helix. The thick and thin blue line at the bottom indicates the position of the folded or unfolded part of the kinked helix, respectively. Orange stars represent the hydrophobic residues at the helix the C-terminal part (C) First principal component. ** p-value vs respective model including Myr < 0.01 (D) RMSD of the kinase domain excluding the *α*_I_ helix. (E) Amplitude of N-C-lobe opening motion (difference between smallest and largest distance between COM of lobes). Centerlines of boxplots denote the mean, box edges the upper and lower quartile. Whiskers represent 1.5 × inter-quartile range. ×× p-value vs whole + Myr < 0.01

**Figure 2.**
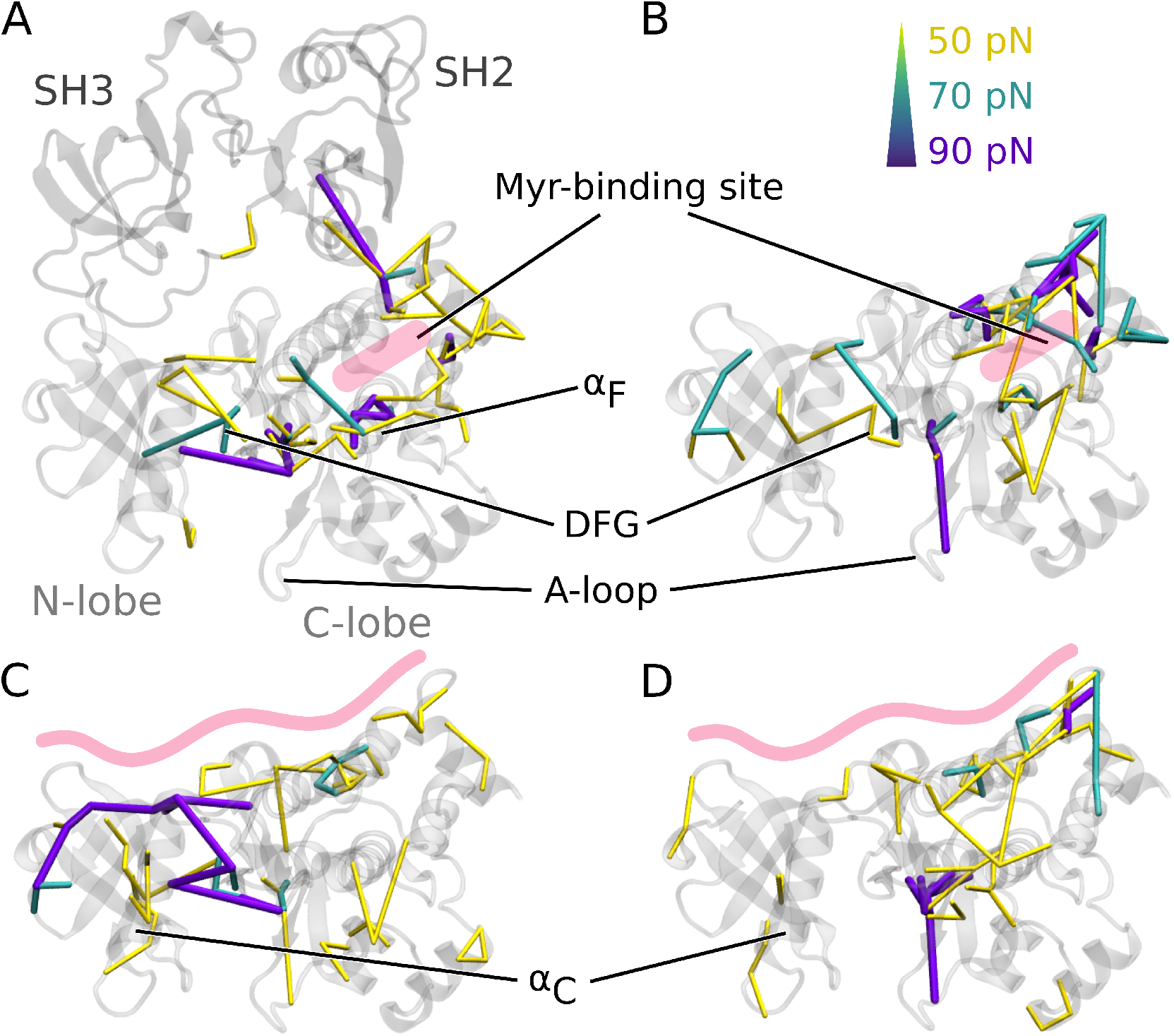
Residue pair-wise force differences. The protein is shown grey. Difference clusters with the indicated force threshold are visualized when at least 3 residues are connected. As indicated in light red, the top row compares differences between simulations with and without Myr in (A) the SH3-SH2-kinase complex or (B) a sole kinase domain with a kinked *α*_I_ helix. The bottom row compares simulations with and without the SH-domains (C) with Myr or (D) without Myr.

The *α*_I_ helix as well as the Myr binding site are located at the C-terminus of the domain. To investigate whether changes on this site of the kinase affect the dynamics of the rest of the domain and thus its activity, we turned to principal component analysis (PCA). For comparison, we simulated the kinase additionally in an active conformation. This model is similar to the model with a straight *α*_I_ helix without Myr, but differs in the conformation of the DFG-motif and *α*_C_ helix, which are both in the ‘in’ conformation. Our analysis shows that the collective motions of the kinase along the first principal component (PC1), which describes an opening and closing of the N- and C-lobe with respect to each other, are closer to that of the active model when Myr was not bound (Fig. 1C, Appendix Fig. 3). This observation is corroborated by a small but significant increase in the RMSD (Fig. 1D) of the kinase and by a larger amplitude of the distance between the N- and C-lobe (Fig. 1E) in the absence of Myr. These observations are counteracted by the presence of the SH-domains, which clamp the kinase in a more rigid state. We note that the differences between the tested conditions are subtle compared to the spread of values within each model. Yet, we observe a significant shift of the conformational ensemble of Abl towards the active state. Thus, Myr unbinding partially activates Abl kinase, while full activation requires the SH-domains to detach.

**Figure 3.**
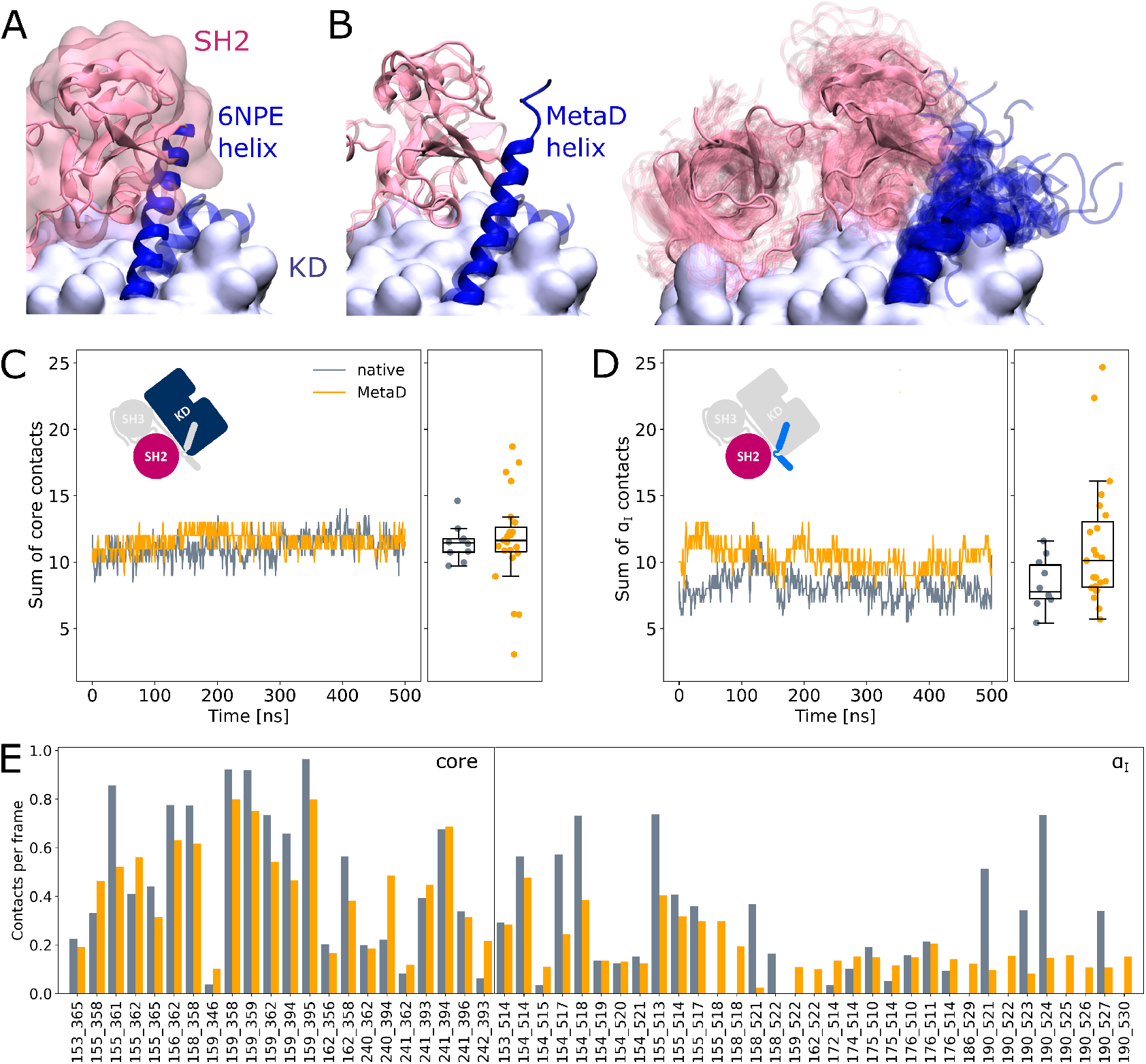
SH2 domain binding is preserved after Metadynamics enhanced *α*_I_ helix straightening. (A) The straight *α*_I_ helix from crystal structures (PDB-ID: 6NPE, blue) clashes with the Abl SH2 domain (light pink). For reference, the structure corresponding to a kinked helix is shown transparently. The kinase domain is shown in opaque surface representation. (B) Helix conformations obtained by Metadynamics simulations are compatible with SH2 binding. One example of a straightened helix is depicted on the left, the right image shows an overlay of all 23 frames with high helicity after simulating them under equilibrium conditions. (C/D) Sum of contacts per frame considering contacts of (C) the kinase domain excluding the *α*_I_ helix or (D) only the *α*_I_ helix. Contacts are considered native when they appeared in our simulations of the Abl SH3-SH2-kinase complex. The ‘MetaD’ contacts were determined from simulations started from Metadynamics frames with high helicity values. (E) Conservation of contacts per frame and residue pair. Contacts, which are present in at least 10% of the simulation time in either native simulations or after Metadynamics are shown. The label ‘core’ denotes contacts excluding the *α*_I_ helix, which start at residue index 505.

### Force distribution analysis reveals collaborative effect of Myr and SH-domains on kinase activity

To understand how Myr or the SH domains influence Abl dynamics and consequently activity we used force distribution analysis (FDA) to decipher allosteric pathways (Fig. 2). In short, we compute time-averaged forces between any pairs of residues as obtained from equilibrium MD simulations and determine the difference between two conditions. Comparing simulations of the SH3-SH2-kinase complex with and without Myr shows high force differences at the active site (Fig. 2A). They are transmitted via the *α*_F_ helix. Residues G402 (DFG-motif), R381 and E305 (*α*_C_ helix) are linked by a 90 pN cluster, which also includes additional residues from the A-loop, DFG-motif and *α*_C_ helix at 50 pN and 70 pN force difference between the Myr bound and unbound state. This likely contributes to *α*_C_ helix outward rotation seen in inactive states of Abl by replacing the E305-K290 interaction stabilizing the inward rotation (***Levinson et al., 2006***; ***Dölker et al., 2014***; ***Liu et al., 2022***). Another high force difference can be seen between the Myr binding site and the SH2-domain, which underlines previous reports that Myr can reinforce SH2-domain docking to the kinase C-lobe (***Nagar et al., 2003***). Performing the same Myr vs. no Myr comparison on simulations of a kinase domain without the SH domains, however, reveals that the allosteric transmission pathway to the active site is missing (Fig. 2B). Additionally, the connections between the A-loop and *α*_C_helix are weakened or gone. The highest differences are mainly visible at the Myr binding site, involving all four helices making up the binding pocket. This reflects the increased flexibility of the second part of the *α*_I_ helix in the absence of Myr (compare Fig. 1B). The only high force difference outside the Myr-binding site comprises residue K419 of the A-loop, which likely contributes to the collapsed A-loop conformation observed in inactive Abl. Overall, FDA results indicate that Myr binding by itself has a long-range allosteric effect on the kinase domain and in particular its active site. This effect is in agreement with the observed decreases in RMSD and N-C-lobe opening amplitude, and goes beyond locally controlling the conformation of the *α*_I_ helix. However, the SH domains, by locking the kinase domain, are required for the inhibitory effect of Myr to be properly transmitted to the active site.

We then went on to focus on the allosteric impact of SH-domain binding to the kinase both in the presence or absence of Myr. In simulations with Myr, medium force differences spread over the whole kinase domain (Fig. 2C), while the highest differences are located at the hinge between the N- and C-lobe and extend to the active site. They connect residues between the A-loop and *α*_C_ helix. In detail, the cluster includes all three residues from the DFG motif (D400, F401, G402) at 70 pN force difference and at 90 pN links R405 and R381 to E305 and E311 from the *α*_C_ helix to maintain its outward rotation. In Abl’s active conformation, R381 forms hydrogen bonds to R405 and pY412 to stabilize the extended conformation of the A-loop (***Xie et al., 2020***). By turning its sidechain towards E305 this interaction is disrupted. Furthermore, the rotation of R381 might contribute to the DFG-flip towards the inactivating out conformation and accompanying disruption of the R-spine, which consists of residues M309, L320, H380 and F401 (***Azam et al., 2008***). ***Levinson et al. (2006***) have proposed the DFG-flip to be facilitated by *α*_C_ helix outward movement since it creates space for rotation of F401. Rotation of R381 additionally pulls the sidechain of the adjacent H380 away from F401, which might aid its rotation towards the active site. In the absence of Myr, we still observe force differences spread over the kinase-SH-domain interface (Fig. 2D). However, all force differences spanning the active site are gone. Forces are instead transmitted to residues of the A-loop, which is located at the opposite site of the kinase domain C-lobe with respect to the SH3-SH2 binding site. Interestingly, this involves the same residue pair K419-D382 as in the Myr bound or unbound comparison without the SH domains, suggesting allosteric pathways to be shared by Myr and the SH-domains. These results further underline that both Myr and the SH domains have an impact on the kinase domain that extends to the A-loop, but neither are sufficient to fully inhibit Abl by themselves and they instead exert a collaborative effect.

### *α*_I_ Helix rearrangements following Myr unbinding can occur without SH domain detachment

It is commonly thought that Myr unbinding induces *α*_I_ helix straightening as the next step in the allosteric activation pathway of Abl (***Nagar et al., 2003***). This is supported by the fact that the straight helix is incompatible with kinase inhibition by SH2 domain binding to the C-lobe of the kinase as observed in crystal structures of Abl (Fig. 3A). In contrast, as mentioned above, we did not observe spontaneous straightening of the *α*_I_ helix in the absence of Myr or the SH-domains. However, the folding process might happen on a timescale longer than accessible with conventional MD simulations. To accelerate the process and asking how *α*_I_ helix straightening impacts SH2 binding, we turned to Metadynamics simulations. We enhanced the sampling of conformations with high helicity (***Pietrucci and Laio, 2009***) of residues 515-521, which correspond to the region interrupting the helical fold of the *α*_I_ helix in its kinked conformation.

We observed that straightening of the helix is possible in presence of the SH domains, with transient straightening events occurring during the Metadynamics simulations without SH domain detachment (example shown in Fig. 3B, left). To asses the stability of the straight conformations without the bias introduced by Metadynamics, we took 23 frames with high helicity across different trajectories, and simulated them using equilibrium conditions for another 500 ns. Residues from M515 onward remained folded in 10 cases, (partially) unfolded in 9 cases and fell back to a conformation resembling the kinked helix in 4 cases (Fig. 3B, right). The kinase-SH2 interface is stable as quantified by the sum of contacts per frame (Fig. 3C,D), and the SH2 domain has to only marginally adjust its position for helix straightening. This was already evident from only transient unbinding of the SH2 domain during a few Metadynamics simulations (Appendix Fig. 4). When considering contacts which are formed in at least 10% of the simulation time, 61 out of 69 (88.4%) contact pairs are conserved compared to MD simulations of the SH3-SH2-kinase complex with a kinked *α*_I_ helix (Fig. 3E). Unsurprisingly, most contact losses are observed between the SH2 domain and the *α*_I_ helix, which changed conformation during the Metadynamics simulation. The average number of native contacts per frame excluding the *α*_I_ helix (’core’ in Fig. 3E) reduces from 0.35 to 0.32 with a mean absolute error of 0.094. We conclude that helix conformational changes are certainly possible without prior unbinding of the SH2 domain.

**Figure 4.**
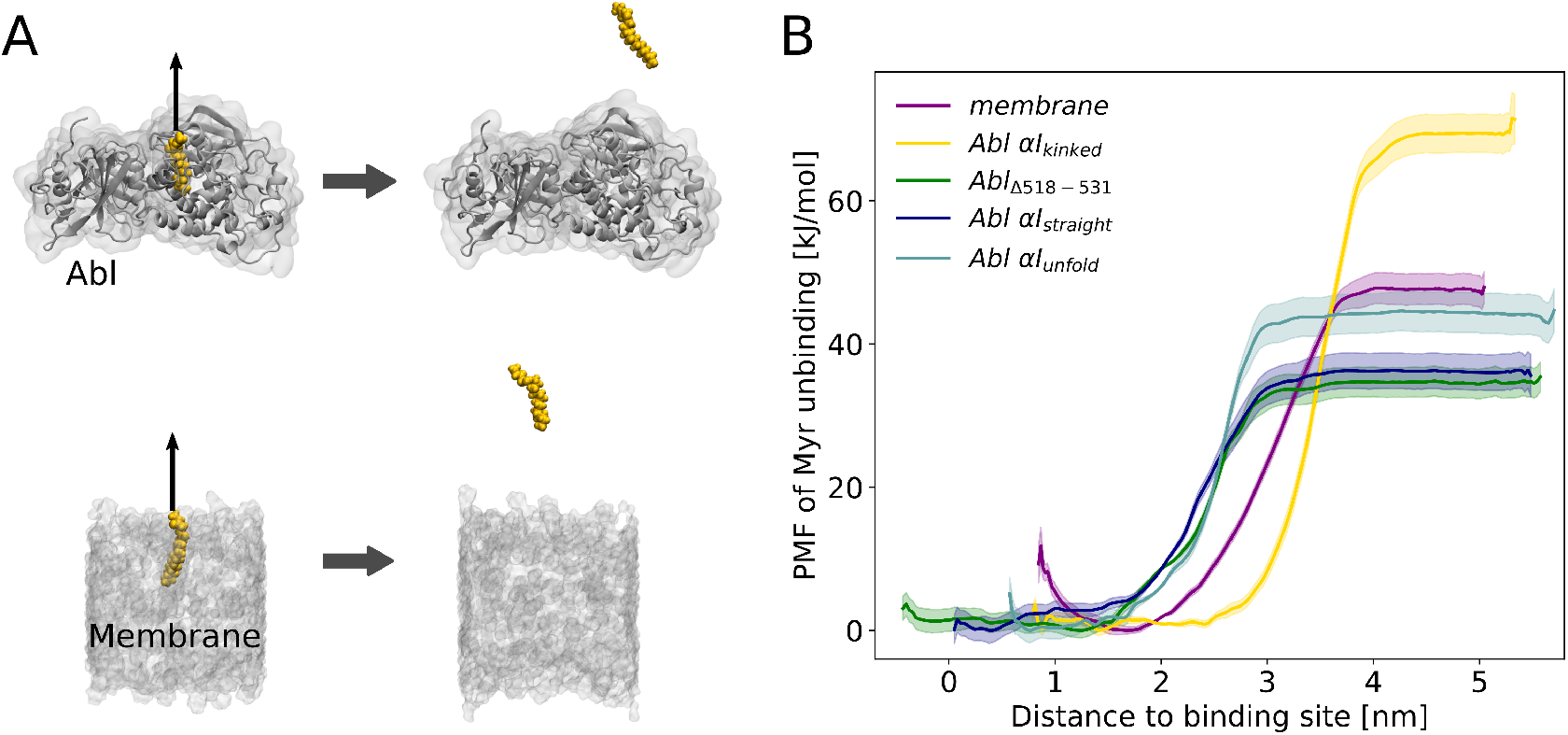
Myr binding to the membrane can compete with Myr binding to Abl. (A) Umbrella Sampling was performed after pulling Myr from its protein binding pocket or a POPC membrane. (B) Potential of mean force (PMF) profiles of Myr unbinding. For comparison, we used an Abl SH3-SH2-kinase complex with a partially unfolded or straight *α*_I_ helix conformation as obtained by Metadynamics simulations, with the residues after the kink (518-531) deleted, or with the kinked conformation.

### Myr can bind to the cellular membrane

Having established how Myr acts as an allosteric effector of Abl, the open question remains where the hydrophobic Myr is stored after detaching from its allosteric binding pocket in the kinase for pre-activation. Conceivable possibilities are either other proteins or lipid membranes. To the best of our knowledge no proteins acting as Myr secretion factors for Abl have been identified. Direct membrane binding of Abl is not widely recognized and the membrane proximal fraction of Abl is not affected by Myr removal (***Hantschel et al., 2003***). However, enrichment of Abl at membranes has been observed upon displacement of Myr by an allosteric inhibitor (***Choi et al., 2009***). We therefore reasoned that Myr stays within its binding pocket due to lack of other binding partners while Abl is localized in the cytoplasm. However, if Abl is recruited to membrane proximal regions by binding to actin filaments with its F-actin binding domain upon cell adhesion (***Woodring et al., 2002***), Myr can insert into these membranes. We set out to explore the thermodynamic probability of this scenario by determining the free energy of Myr unbinding from either a lipid bilayer or its protein binding pocket using Umbrella Sampling simulations (Fig. 4).

Displacing Myr from a simple POPC membrane involves a free energy of unbinding, or Potential of Mean Force (PMF), of 45.6 ± 2.7 kJ/mol. Assessing the free energy of Myr unbinding from Abl kinase by Umbrella sampling for comparison is less straightforward due to the involved conformational change of the *α*_I_ helix. In equilibrium MD simulations, we observed the kinked *α*_I_ helix conformation to be destabilized if Myr is absent (Fig. 1B). X-ray structures with an empty Myr pocket support this, and suggest helicity to prevail until residue ∼519, while more C-terminal residues are largely disordered (Appendix Fig. 2). To determine the free energies for Myr unbinding from Abl kinase that reflect this conformational propensity, we carried out Umbrella Sampling simulations using an Abl model with a partially unfolded *α*_I_ helix obtained during Metadynamics simulations. In addition, as extreme cases, a truncated version in which residues 518-531 are absent, and a straightened helix conformation were used. For all of these models, the free energy of Myr unbinding from the protein is similar or smaller compared to unbinding from the membrane, supporting the notion that Myr binding to the membrane is thermodynamically possible. This result is also supported by previous measurements. The dissociation constant K_D_ for binding of a myristoylated peptide from Abl’s kinase domain has been determined to be 2.3 *µ*M (***Hantschel et al., 2003***), corresponding to a binding free energy of 32.4 kJ/mol (Table 1), in good agreement with the calculated free energies using Abl models with a truncated or straight *α*_I_ helix conformation. In contrast, the free energy for Myr unbinding from Abl was significantly higher when using an Abl model based on the crystal structure with Myr bound (i.e. with a kinked *α*_I_ helix). This can be explained by the hydrophobic residues lining the Myr binding pocket, which interact with Myr and are solvent exposed after unbinding.

**Table 1.**
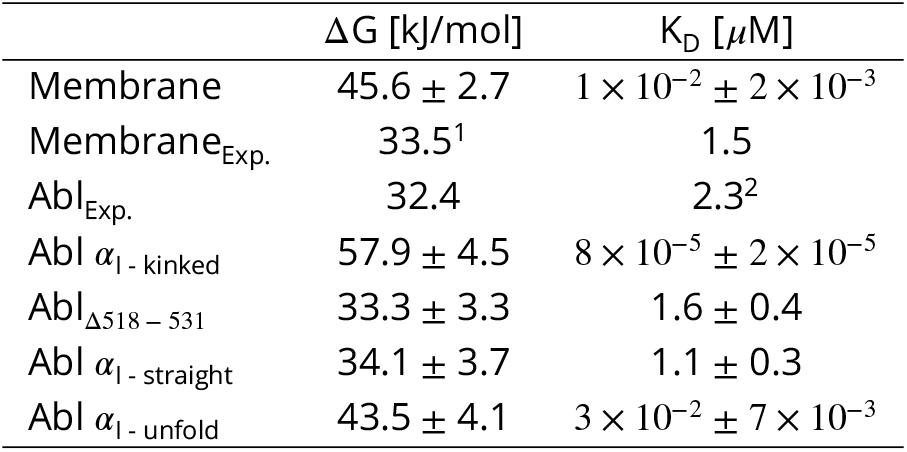
ΔG values derived from Umbrella Sampling simulations or observed experimentally (^1^***Peitzsch and McLaughlin (1993***), ^2^***Hantschel et al. (2003***)). K_D_ values are calculated using the Nernst equation with the assumption of an equal number of binding sites of 1 per Myr. Errors in the ΔG values were obtained by bootstrapping and propagated for K_D_ calculation.

Overall, our Umbrella Sampling results indicate that indeed helix rearrangements have to be taken into account to reflect the correct energy of unbinding. We note that these helix rearrangements of course contribute to the actual free energy of Myr unbinding but have not been covered during the limited timescale of Umbrella Sampling. The comparison to the experimental value, however, suggests that this approximation is feasible. We note that the free energy of Myr unbinding from a membrane obtained from simulations is higher than the value of 33.5 kJ/mol that has previously been determined experimentally (Table 1) (***Peitzsch and McLaughlin, 1993***). Notwith-standing, the two experimental studies of Myr-Abl binding and Myr-membrane insertion confirm our major result that the involved free energies are comparable. Thus, we conclude that insertion of a single myristoyl moiety into a lipid bilayer can thermodynamically compete with binding of Myr into the known pocket of Abl’s kinase domain.

## Discussion

In this study we have used equilibrium and enhanced sampling simulation techniques to understand Abl’s allosteric inhibition by Myr and the SH domains. We were not only able to describe the effect of different *α*_I_ helix conformations as well as Myr and SH domain binding on the overall dynamics of Abl’s kinase domain, but could also explain the observations by identifying the pathways from the allosteric site to the active center at residue level resolution. We observed that both Myr and the SH domains by themselves were able to impact residues at the A-loop. However, they act in concert for full inhibition: We saw how forces are transmitted from the Myr binding site to the active site along the *α*_F_ helix only in the presence of the SH domains. Vice versa, we observed how the SH domains act on the hinge between N- and C-lobe only when Myr was bound to the kinase domain, affecting known hallmarks of Abl kinase activity such as the *α*_C_ helix, the DFG-motif and the R-spine (Fig. 2). This is in line with and explains at molecular detail that a kinase domain bound to Myr or the allosteric inhibitors GNF-2 and GNF-5 without the SH domains or Abl mutants deficient in Myr binding display increased activity (***Choi et al., 2009***; ***Fabbro et al., 2010***; ***Hantschel et al., 2003***).

It has so far remained unclear how exactly Abl activation progresses upon Myr unbinding. Using equilibrium and Metadynamics simulations, we were able to show that the *α*_I_ helix gains flexibility upon Myr unbinding and that both straightening and partial unfolding of the *α*_I_ helix are possible even without prior unbinding of the SH domains. Both types of helix rearrangements are compatible with crystal structures of Abl’s kinase domain (Appendix Fig. 2). The helix fold continues longer compared to the kinked conformation seen in crystal structures of the Myr-bound SH3-SH2-kinase complex (***Nagar et al., 2003***), but is often not resolved far beyond the position of the kink, in agreement with the high flexibility of this most C-terminal part we observe in our simulations. The helix rearrangements upon Myr unbinding involve an only slight adaptation of the SH2 position as evidenced by shifts in residue pair contact counts (Fig. 3C). This observation is in line with SAXS and solution NMR data showing that an Abl mutant incompetent in Myr binding or apo-Abl remain in the assembled conformation (***Badger et al., 2016***; ***Skora et al., 2013***). We suggest that the different *α*_I_ helix conformations and/or binding poses of the SH2 domain explain why apo-Abl has failed to crystallize so far. The observation of an allosteric effect of Myr unbinding leading to a pre-activated state with an altered SH-kinase interface implies that factors in addition to Myr unbinding contribute to full Abl activation. Transient unbinding events of the SH domains, as we also observed in a few Metadynamics simulations, could enable phosphorylation of Y245 on the SH2-kinase linker (***Brasher and Van Etten, 2000***) or binding of activator substrates to block rebinding of the SH domains (***Wang, 2014***), thereby advancing Abl activation.

It is widely accepted that Myr unbinding leads to Abl activation. However, the non-myristoylated Abl isoform 1a, which only differs from the 1b isoform by the N-terminal half of the N-cap, is not de-regulated (***Van Etten, 2003***; ***Nagar et al., 2003***). This suggests that additional factors contribute to Abl activation since loss of Myr can apparently be compensated. We addressed the so far unanswered question where Myr binds to after it has left its protein binding pocket. The fatty acid has to be stored in a hydrophobic location or it will quickly rebind to its binding pocket and switch Abl back into its inactive state. We here propose the membrane as an anchoring point for Myr, identifying a new role for Abl localization and regulation (Fig. 5), which is in line with the fact that protein myristoylation is usually involved in membrane recruitment (***Resh, 2016***). By harbouring Myr, the membrane would stabilize the pre-activated state, allowing time for activating phosphorylations of Y412 on the A-loop or Y245 (***Brasher and Van Etten, 2000***), permanent SH domain detachment, or SH2 domain binding to the kinase N-lobe. All of these events enhance kinase domain transitions to the fully active state. Our results show that if rearrangements of the *α*_I_ helix are taken into consideration, membrane binding is energetically as favorable or potentially even more favorable than protein binding, a tendency also found when comparing experimental measurements (Table 1) (***Hantschel et al., 2003***; ***Peitzsch and McLaughlin, 1993***). Previous reports that Abl localizes to membranes upon Myr displacement from the protein by an allosteric inhibitor (***Choi et al., 2009***) or upon kinase domain deletion (***de Oliveira et al., 2013***) confirm that Abl is indeed capable of binding to membranes and that the membrane competes with the protein hydrophobic pocket for Myr binding. However, ***Choi et al. (2009***) revealed enrichment of Abl at both endoplasmic reticulum membranes and the outer cell membrane, indicating that Myr attached unspecifically to the hydrophobic environment of any membrane.

**Figure 5.**
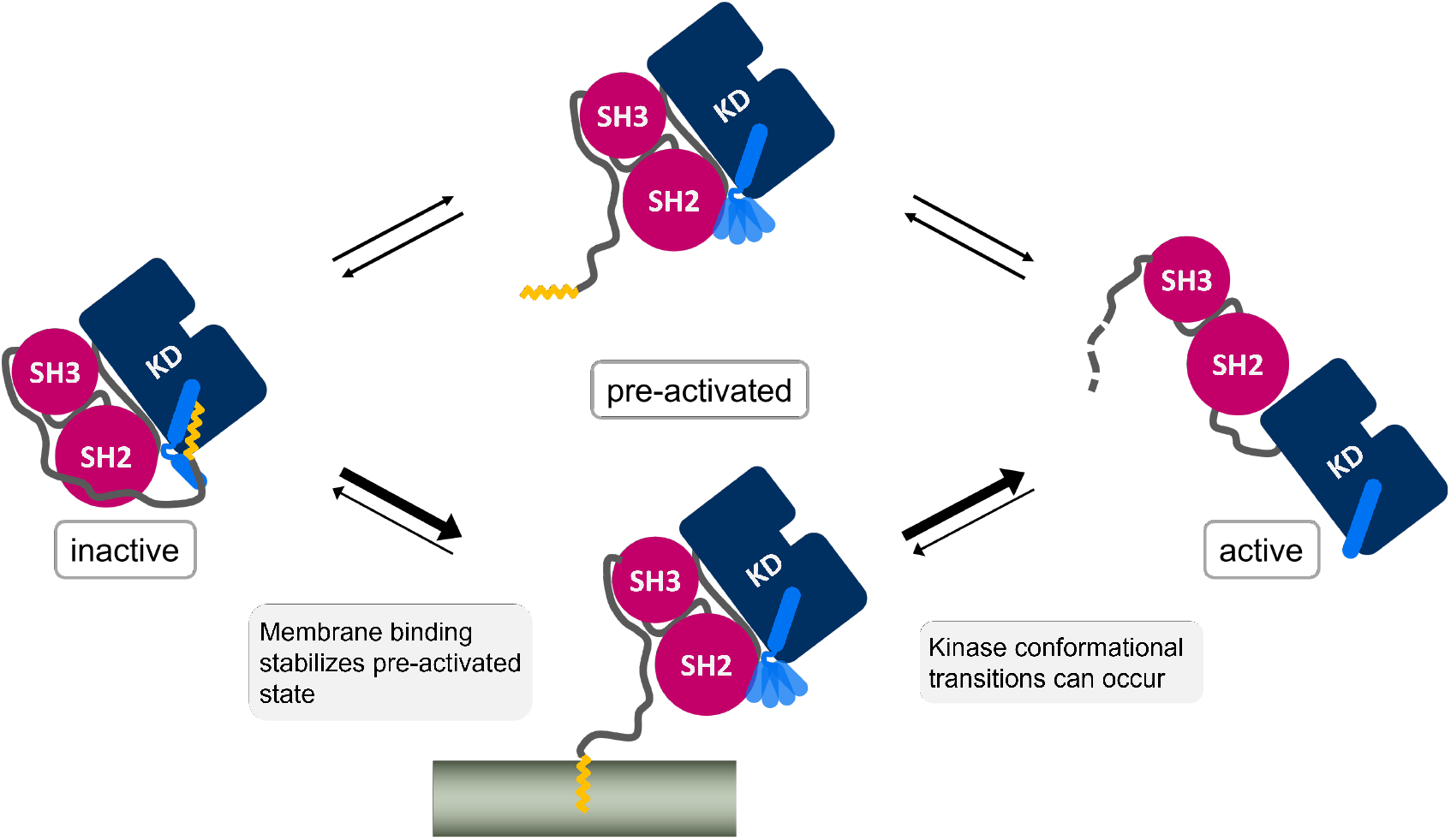
Membrane binding of Myr enhances Abl activation by stabilizing the pre-activated state.

The high free energy difference we determined for Myr unbinding from the SH3-SH2-kinase model with a kinked *α*_I_ helix is consistent with Abl being well soluble in aqueous solution such as the cytosol (***Nagar et al., 2003***), which prevents unspecific binding. At the same time, it explains why the membrane proximal fraction of Abl is not decreased by Myr deletion (***Hantschel et al., 2003***). If Abl gets recruited to regions close to cellular membranes by binding to the cytoskeleton with its C-terminal F-actin binding domain, the free energy difference that has to be overcome reduces from ∼58 kJ/mol to ∼12 kJ/mol (ΔΔG between Myr unbinding from Abl *α*_I - kinked_ and membrane, Table 1). The difference is further balanced by helix rearrangements occurring simultaneously or subsequently to Myr unbinding from Abl. This agrees well with the fact that Abl gets activated after localization to focal contacts (***Lewis et al., 1996***; ***Woodring et al., 2002***). Membrane binding could therefore be considered an extra regulatory layer, which can be carefully balanced by localization cues or membrane composition. In fact, N-cap residues 15-60, which have been shown to be irrelevant for Abl inhibition (***Hantschel et al., 2003***), encompass a number of basic residues (K24, K28, K29, R33) and could interact with acidic membrane lipids found at focal contacts. Src kinase also features basic residues near its myristoylation site which stabilize the interaction with PIP_2_-rich membranes (***Daday et al., 2022***). We speculate that a membrane also enhances Myr unbinding from Abl and Myr insertion by using these residues as a guide for Myr.

In summary, we have shown that Myr and the SH domains act in concert to inhibit Abl kinase activity and visualized the detailed allosteric transmission pathway from the Myr binding site to the active site. We propose a unique dual role of Myr for Abl: in addition of Abl inhibition it also localizes Abl to the membrane. Proximal membranes accommodating Myr would thus aid the allosteric activation of Abl by stabilizing its pre-activated state. This novel crosstalk can be directly tested by biochemical or cell experiments, and is ideally suited to be exploited by new therapeutic approaches that allosterically target Abl.

## Methods

### Modelling of Abl structures for simulations

We based our Abl models on a crystal structure including the SH3, SH2 and kinase domains as well as part of the N-Cap and Myr (PDB-ID: 2FO0). This structure has a D355N mutation, which we reversed using Modeller (***Sali and Blundell, 1993***). Furthermore it is bound by the inhibitor PD-166326 (6-(2,6-dichlorophenyl)-2-[3-(hydroxymethyl)phenyl]amino-8-methylpyrido[2,3-D]pyrimidin-7(8H)-one). Since we aimed for a more physiological structure we replaced this inhibitor by ATP by doing the following modifications: We overlayed the kinase domain with an ATP-analog-bound Abl structure (PDB-ID: 2G2F, RMSD of 1.7 Åacross all atoms). We converted the ATP-analog 112 (thiophosphoric acid O-((adenosyl-phospho)phospho)-S-acetamidyl-diester) to ATP by deleting the acetamidyl group and changing the sulfur-atom to oxygen. The A-loop of 2FO0 was pushed upwards by the inhibitor and the ATP-positioning loop clashed with the ATP-phosphates. We therefore refined these two loops based on homology with the 2G2F structure. We modelled the structures with a straight *α*_I_ helix by creating a homology model of the modelled ATP-bound kinase and PDB structure 6NPE (RMSD 1.47 Å) using Swiss model (***Waterhouse et al., 2018***). For the active kinase domain model, we used a structure with the A-loop in DFG-in conformation that did not have an inhibitor bound to it (PDB-ID: 2G2I) and filled in missing P-loop residues using Modeller.

### Equilibrium MD

We used Gromacs (***Van Der Spoel et al., 2005***) version 2018.5 and 2020.3 along with the CHARMM36 force field (March 2019 version) (***Best et al., 2012***; ***Huang and MacKerell, 2013***; ***Klauda et al., 2010***). The proteins were placed in a dodecahedral simulation box with distance of 2 nm between the box boundary and nearest protein atom. The box was filled with TIP3P water and 150 mM NaCl to neutralize charges. For energy minimization, we applied the steepest descent algorithm with a step size of 0.1 nm and a force tolerance of 1000 kJ/mol. During the two equilibration steps, the positions of the peptide backbone heavy atoms were restrained with a force constant of 1000 kJ/mol/nm. The temperature was equilibrated to 300 K using the v-rescale thermostat (***Bussi et al., 2007***) with a coupling constant of 0.5 ps in a 100 ps simulation in the NVT ensemble. This was followed by a 1 ns simulation in the NPT ensemble to adjust the pressure to 1 bar using isotropic pressure coupling with the Berendsen barostat (***Berendsen et al., 1984***) with a coupling constant of 5 ps. After equilibration, we switched to the Parinello-Rahman barostat (***Parrinello and Rahman, 1981***) for pressure coupling. For temperature coupling, we kept the v-rescale thermostat or switched to the Nosè-Hoover thermostat (***Nosé, 1984***; ***Hoover, 1985***). We simulated 10 replicates for 500 ns for each Abl model and thermostat. Since the sets of simulations with different thermostats show the same trends (Appendix Fig. 5), we decided to combine their analyses. For integrating the equations of motion, the leap-frog integrator with a time step of 2 fs was used. All bonds involving hydrogen atoms were constrained using the LINCS algorithm (***Hess et al., 1997***). Van-der-Waals interactions were smoothly switched to zero between 1.0 and 1.2 nm using the force-switch method. Long range coulomb interactions beyond 1.2 nm were treated with the PME method (***Darden et al., 1993***).

### Force Distribution Analysis

For force distribution analysis (FDA), we concatenated the replicates of each model into a single trajectory and calculated the residue-based pairwise forces (***Costescu and Gräter, 2013***). We included all bonded and nonbonded interaction types and averaged the forces over the whole trajectory. We determined the force-differences between a) simulations with and without Myr using both kinase only (kinked *α*_I_ helix conformation) and SH3-SH2-kinase models and b) simulations of kinase domain only versus SH3-SH2-kinase both in the absence and presence of Myr. We visualized force-difference clusters with a minimum size of 3 residue pairs and a force threshold ranging between 20-100 pN using the fda_graph tool and VMD (***Humphrey et al., 1996***).

### Metadynamics

We used Gromacs 2018 patched with Plumed version 2.5.2 (***Bonomi et al., 2009***; ***Tribello et al., 2014***) to perform the Metadynamics simulations. As the collective variable (CV) we chose the so called *alpharmsd*, which compares backbone distances between adjacent amino acids to typical helix distances and thereby functions as a measure for helicity (***Pietrucci and Laio, 2009***). We performed the simulations on our SH3-SH2-kinase complex model without Myr and applied the bias to residues 515-521, which correspond to the unfolded region in the kinked *α*_I_ helix. We tested bias factors between 10 and 100, of which 20-30 gave the best results. Gaussians with an initial height of 1.2 kJ/mol and sigma of 0.02 and 0.03 were added every 500 integration time steps. The value of the CV and the bias potential were printed every 100 steps. Transient straightening events occured in 11 out of 14 simulations. After straightening the helix often partially unfolded, which is a consequence of Metadynamics penalising the already visited conformations. We extracted a total of 23 frames with high alpharmsd from each Metadynamics run and simulated them for 500 ns as described above in equilibrium conditions.

### Umbrella Sampling

We built a pure POPC membrane and embedded a myristoylated Gly residue such that the fatty acid chain is parallel to the membrane lipids and Gly faces the lipid head groups using Charmm-Gui (***Jo et al., 2008***; ***Lee et al., 2016***). We considered SH3-SH2-kinase Abl models with a kinked, truncated, straight or partially unfolded *α*_I_ helix conformation. For the truncated *α*_I_ helix we deleted residues 518-531. Models with a straight or unfolded *α*_I_ helix were obtained from the Metadynamics simulations. We used frames from the end of the 500 ns equilibrium simulations as start frames for pulling simulations. Pulling and Umbrella Sampling was done similarly as reported previously (***Lemkul and Bevan, 2010***). We created a triclinic box with at least 2 nm distance between the protein and the box boundaries and elongated the box to 12 nm to provided space for pulling out Myr. The dimension for the membrane system were approximately 5 × 5 nm along the membrane plane and extended to 11 nm in the pulling direction vertical to the membrane plane. We used Gromacs 2020 for the Umbrella Sampling simulations. The systems were solvated, neutralized, energy minimized and equilibrated as described in the Equilibrium MD section. We pulled on the heavy atoms of the C-terminal NME cap of the Myr-Gly residue at 0.1 and 0.05 m/s with a spring constant of 500 kJ/mol/nm^2^. Per model and pulling velocity we performed at least 3 replicates. Reference groups for pulling were the lipids of the upper membrane leaflet within a 1.5 nm radius around Myr, or the protein backbone residues of the four *α* helices comprising Abl’s Myr binding site.

To obtain the starting configurations for the umbrella sampling windows, we extracted frames from the pulling trajectories with a spacing along the COM distance between pulling groups of 0.1 nm for the first 1 nm and 0.2 nm beyond that. This allows to sample the initial unbinding stage at more detail. The fully unbound state is sufficiently covered at a distance of approximately 4.5 nm as evidenced by the plateau in the PMF plots. We simulated each window for 50 ns. For analysis with the weighted histogram analysis method (WHAM) (***Kumar et al., 1992***), we discarded the first 10 ns simulation time of each window. We estimated the error using bootstrapping with the b-hist method included in gmx wham (***Hub et al., 2010***) with 100 bootstraps.

## Data Availability

Source data for all figures has been deposited on the Dryad Digital Repository under the DOI 10.5061/dryad.9cnp5hqnx

## Acknowledgments

We acknowledge funding through the Deutsche Forschungsgemeinschaft (DFG, German Research Foundation) under Germany’s Excellence Strategy – 2082/1 – 390761711, the Klaus Tschira Foundation, the state of Baden-Württemberg through bwHPC, as well as the DFG through grant INST 35/1134-1 FUGG. S.d.B. thanks the Carl Zeiss Foundation for financial support.

## Appendix 1

**Appendix 1—figure 1.**
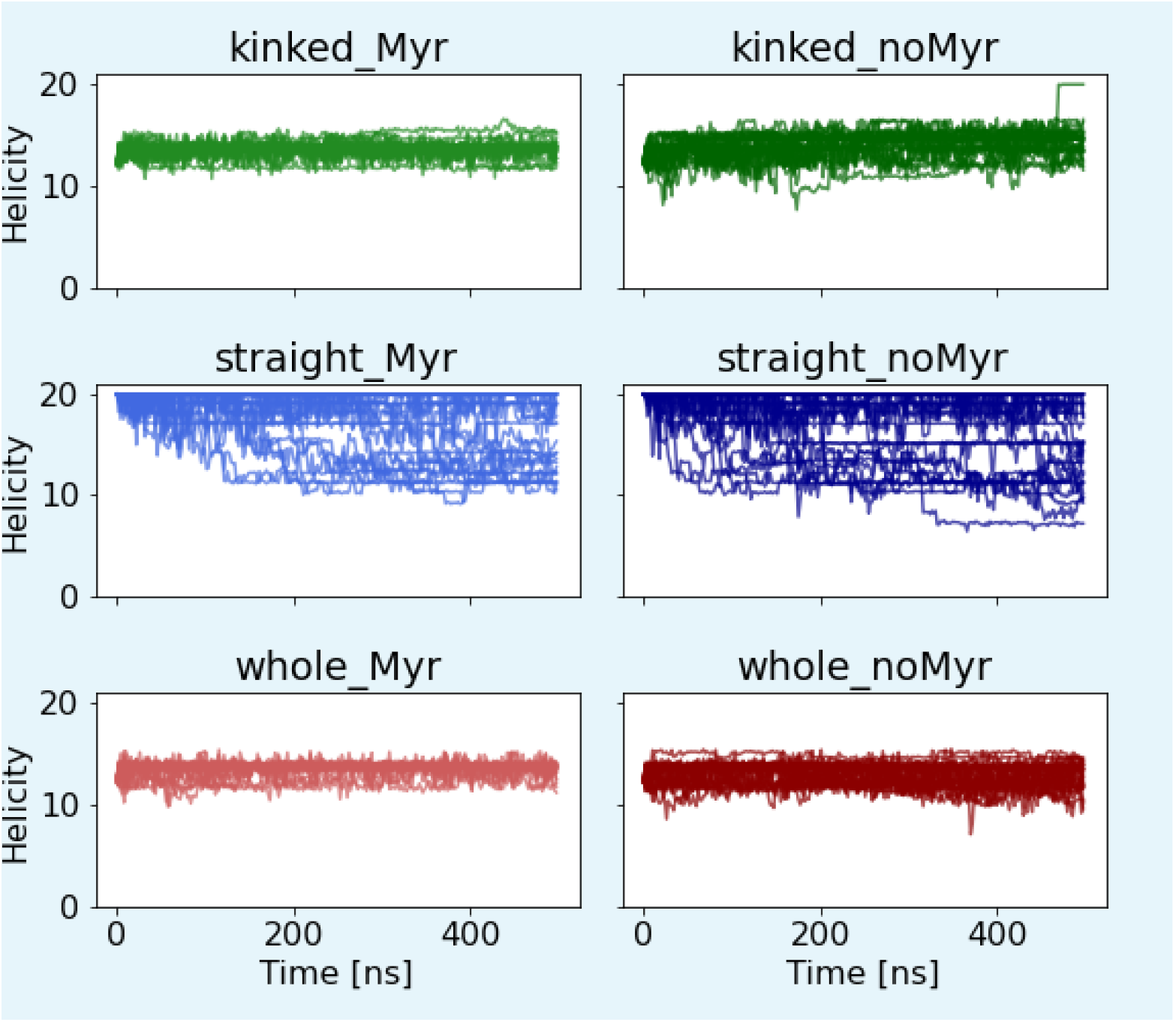
Helicity of the *α*_I_ helix. Helicity is calculated using Plumed’s alpharmsd CV. Straight *α*_I_ helices partially unravel regardless of whether Myr is bound or not. The kinked helices remain stable in all cases albeit with slightly larger helicity fluctuations in the absence of Myr.

**Appendix 1—figure 2.**
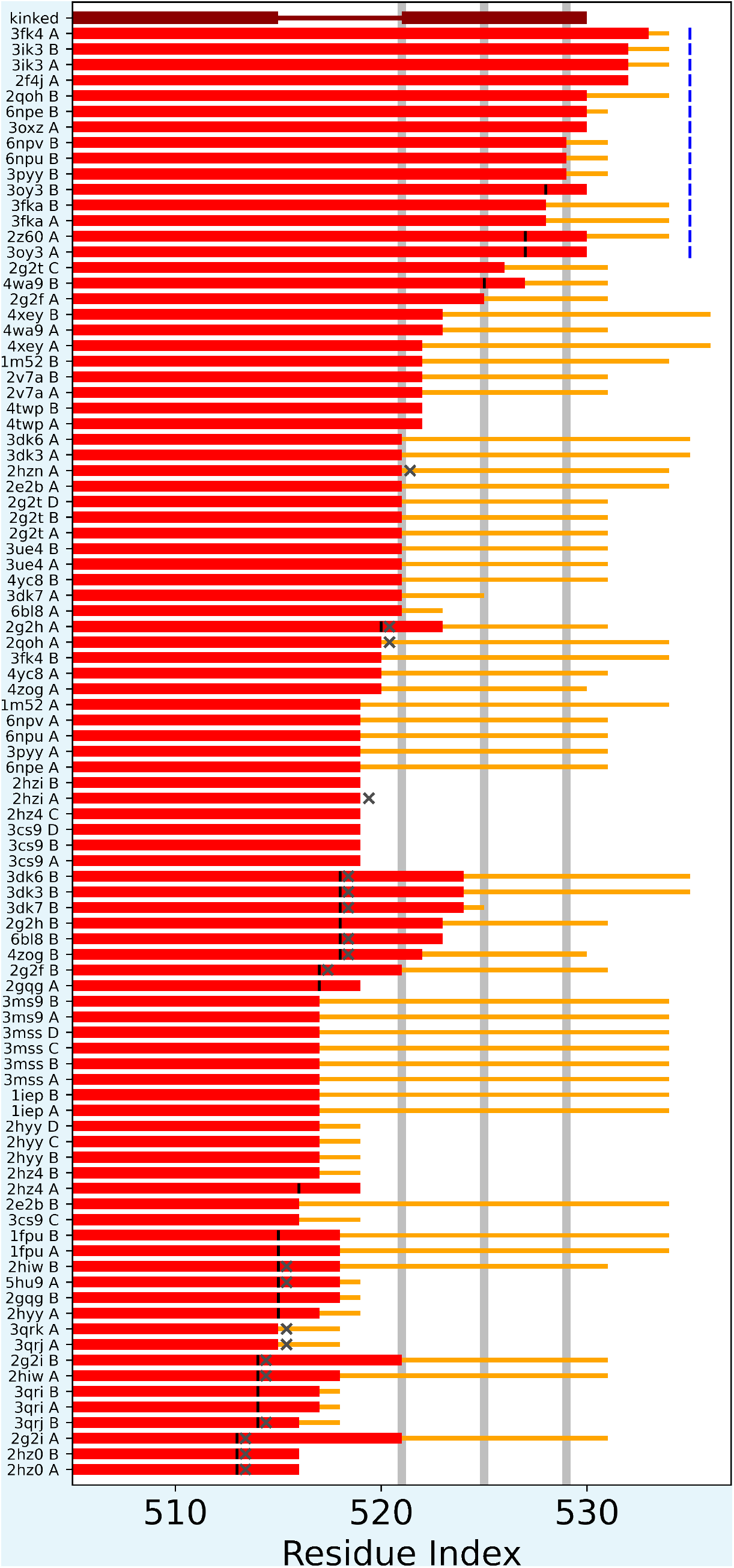
Overview over *α*_I_ helix lengths in Abl crystal structures published on the protein data bank. The kinked *α*_I_ helix structure is shown for reference in dark red. The two thick line segments indicate the folded part, which is interrupted as shown by the thinner line segment. Light grey horizontal lines represent the position of the hydrophobic residues I521, L525 and V529. Light red lines depict resolved residues. Yellow lines represent residues that were present in the construct used for crystallization, but were not resolved. Vertical black lines indicate that the helices where not folded beyond this residue position, dark grey Xs that helix continuation was blocked by another copy of Abl in the crystal lattice. Blue labels illustrate that the straight helix conformation, especially the hydrophobic residues, where protected by another protein copy in the crystal lattice. It can be seen that the majority of helices that continue folded beyond the first hydrophobic residues are stabilized by another protein copy, while helices, which end much sooner are either blocked or the respective residues where not included in the construct used for crystallization.

**Appendix 1—figure 3.**
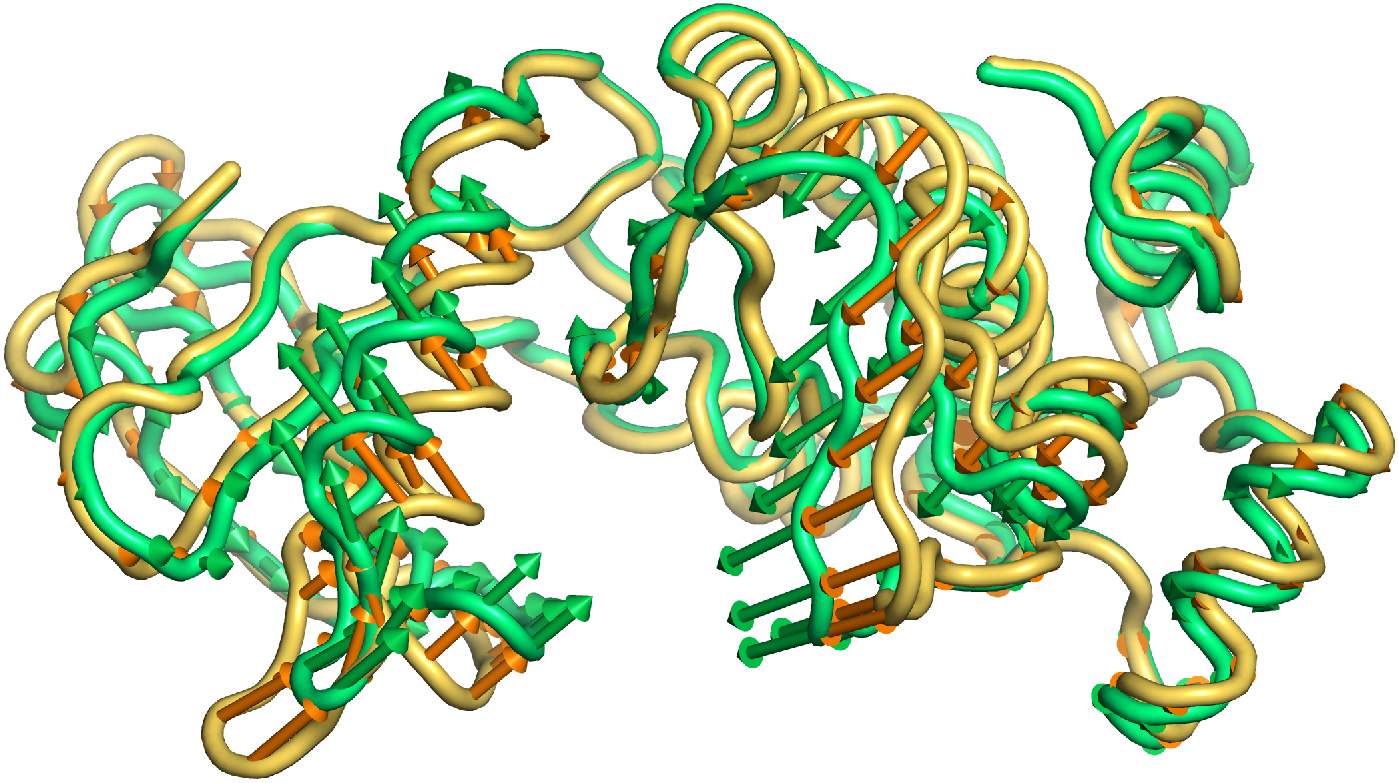
Porcupine plot illustrating the motion described by the first principal component. Orange tones represent the kinase domain in an active conformation, green tones represent the kinase with a kinked *α*_I_ helix bound with Myr. Visualized using PyMol v2.4.1 and the modevectors.py script (http://www.pymolwiki.org/index.php/Modevectors)

**Appendix 1—figure 4.**
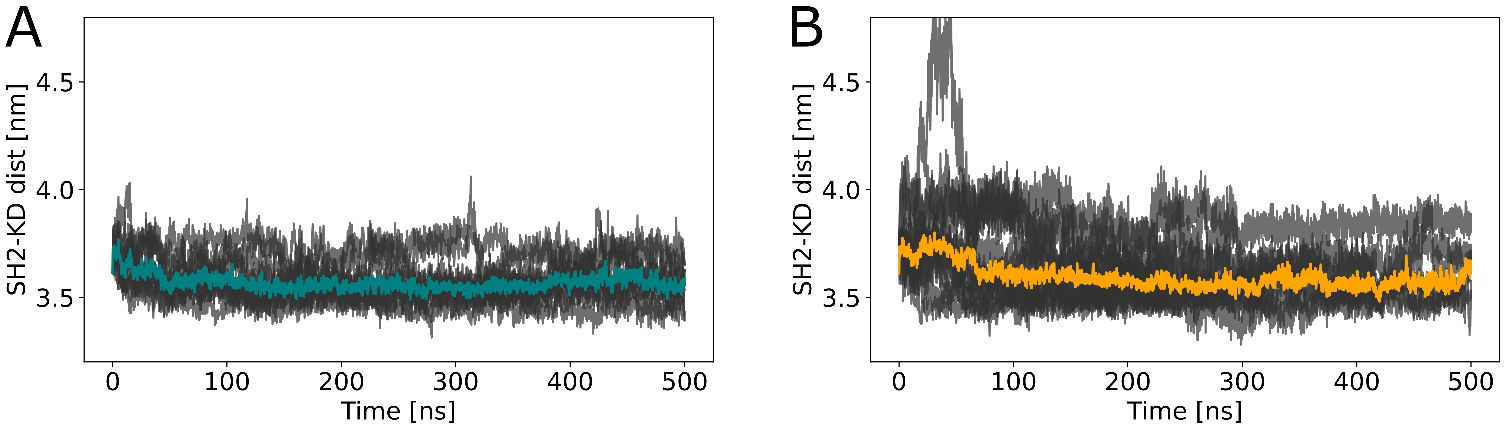
Distances between the center of mass of SH2 and kinase domain during (A) equilibrium MD and (B) Metadynamics simulations. Despite transient unbinding events characterized by distance increases towards the beginning of few Metadynamics simulations, the median distance (depicted as colored lines) is very similar in both cases: 3.56 for equilibrium MD and 3.55 for Metadynamics simulations.

**Appendix 1—figure 5.**
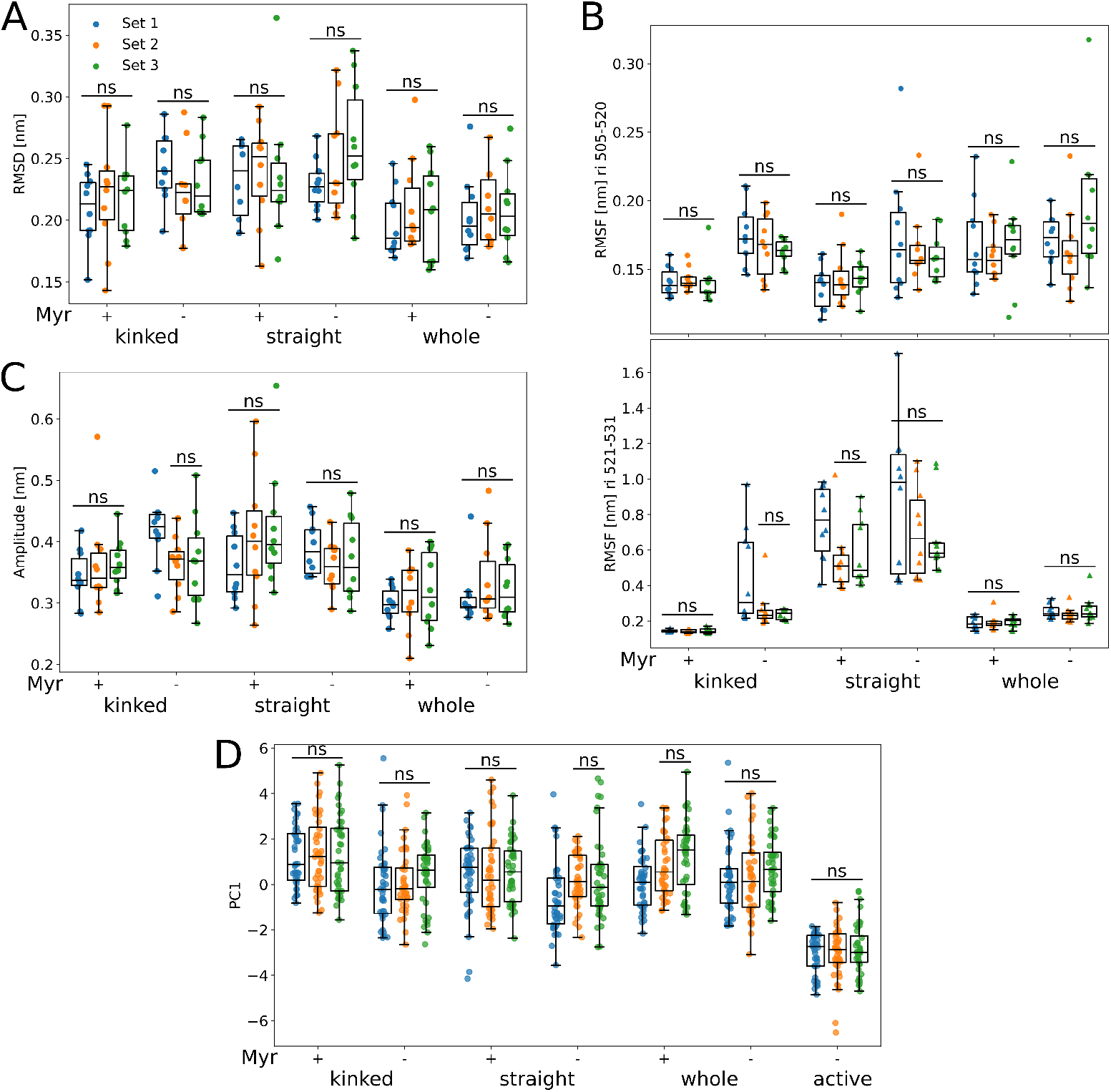
Comparison between different simulation conditions (used to create Figs. 1+2). Set 1 was simulated using Gromacs 2018.5 and the Nosè-Hoover thermostat. The proteins were not capped at the termini. Set 2 and 3 were both simulated using Gromacs 2020.3 and the proteins were capped to remove artificial charges introduced by a premature ending of protein structures, i.e most of the N-terminal linker and the C-terminal disordered region and F-actin binding domain were not part of any of the simulated systems. The difference between set 2 and 3 is the use of the v-rescale and Nosè-Hoover thermostat, respectively. The remaining simulation parameters were as described in the methods part of the main article.

## Notes

### Competing Interest Statement

The authors have declared no competing interest.

